# Dynamic RNA Polymerase II Recruitment Drives Differentiation of the Intestine under the direction of HNF4

**DOI:** 10.1101/2023.11.08.566322

**Authors:** Kiranmayi Vemuri, Sneha Kumar, Lei Chen, Michael P. Verzi

## Abstract

Terminal differentiation requires a massive restructuring of the transcriptome. During intestinal differentiation, the expression patterns of nearly 4000 genes are altered as cells transition from progenitor cells in crypts to differentiated cells in villi. We identified dynamic recruitment of RNA Polymerase II (Pol II) to gene promoters as the primary driver of transcriptomic shifts during intestinal differentiation *in vivo*. Changes in enhancer-promoter looping interactions accompany dynamic Pol II recruitment and are dependent upon HNF4, a pro-differentiation transcription factor. Using genetic loss-of-function, ChIP-seq and IP mass spectrometry, we demonstrate that HNF4 collaborates with chromatin remodelers and loop-stabilizing proteins and facilitates Pol II recruitment at hundreds of genes pivotal to differentiation. We also explore alternate mechanisms which drive differentiation gene expression and find pause-release of Pol II and post-transcriptional mRNA stability regulate smaller subsets of differentially expressed genes. These studies provide insights into the mechanisms of differentiation in a renewing adult tissue.

**HIGHLIGHTS:**

- Dynamic recruitment of Pol II largely drives the vast transcriptomic changes seen during differentiation of mouse intestinal epithelium.
- Smaller groups of differentiated genes are subject to regulation through Pol II pause-release and post-transcriptional mechanisms such as differences in mRNA stability.
- IP-mass spectrometry analysis identifies the first interactome of HNF4 in the differentiated small intestine, finding interactions with chromatin looping and chromatin remodeling proteins.
- HNF4 transcription factors play a critical role in recruiting Pol II to the promoters of essential intestinal differentiation genes.

## INTRODUCTION

Transcriptomic shifts are a fundamental prerequisite for cellular differentiation across various tissues and organs. These shifts involve a dynamic reprogramming of gene regulatory patterns, where specific sets of genes are activated or suppressed to guide a cell’s transformation from a multipotent state into a highly specialized and functional cell type. The intestinal epithelium is a prime example of rapid and perpetual differentiation. In the mucosa of the small intestine, stem cells and actively dividing progenitor cells are predominantly located within the crypts of Lieberkühn.^1^ In contrast, differentiated, non-dividing cells responsible for vital absorptive and secretory functions are found along the protruding villi. The entire lifespan of a stem cell from the base of the crypt until it differentiates as it migrates to the tip of the villus is 3-5 days.^2,3^ Differentiation of the intestinal epithelium from crypts onto villi underlies many areas of intestinal health, encompassing everything from nutrient absorption to immune defense.^4–6^

A fundamental aspect of crypt-villus differentiation lies in the precise control of gene expression, ensuring that cells along this axis maintain distinct identities and functions. Studies centering on *cis*-regulatory elements have provided valuable insights into chromatin patterns and gene regulatory events occurring during differentiation. The coordination between transcription factors CDX2 and HNF4α at enhancer sites helps in maintaining an open chromatin state in differentiated villus cells.^7^ Profiling of H3K27ac, H3K4me2, and DNaseI in crypt and villus cell populations revealed that this state of accessible chromatin is universal across both cell types, which facilitates lineage-specification of differentiated cell types based on the available set of transcription factors.^8^ The upregulation of differentiation specific genes such as those associated with enterocyte metabolism also corresponds with a loss of DNA methylation from the stem to the differentiated cell state along with an increase in enhancer activity, and CDX2 and HNF4 binding.^7,9,10^ This cascade of epigenomic events culminates in a dynamic transcriptome as cells progress from crypts to villi, resulting in the differential regulation of nearly 4000 genes.^11–13^ Nevertheless, it remains unclear how upstream regulators intersect with basic *trans*-regulatory mechanisms, such as the recruitment and regulation of RNA polymerase II at gene promoters.

Surprisingly, the genomic distribution of Pol II along the crypt-villus axis remains unexplored, underscoring the gap in our knowledge of gene regulation during intestinal differentiation. The recruitment of Pol II to target gene promoters is intricately linked to three-dimensional intrachromosomal contacts within topologically associated domains.^14,15^ The spatial organization of chromatin loops brings distal enhancers into close proximity with gene promoters, facilitating the recruitment of transcription factors and the assembly of the transcriptional machinery, including Pol II.^16^ In the context of the intestine, the pro-differentiation factor HNF4 exerts a significant influence on this process, as evidenced by loss of enhancer-promoter looping events (measured by H3K4me3 HiChIP-seq) in villus-specific genes when HNF4 is depleted.^12^

Within the Pol II transcription process itself, multiple regulatory checkpoints influence gene expression through a variety of factors and mechanisms. At the initiation checkpoint, the assembly of the transcription initiation complex and the recruitment of Pol II at the gene promoter mark the commencement of gene transcription. Variations in Pol II recruitment to the gene promoter can lead to substantial alterations in gene expression.^17^ Additionally, promoter-proximal pausing of Pol II (hereafter referred to as ‘Pol II pausing’) has emerged as a prominent regulatory mechanism influencing gene expression.^18–21^ This phenomenon involves the temporary halting of Pol II shortly after transcription initiation, poised to rapidly respond to environmental cues. The dynamic equilibrium between paused and actively elongating Pol II has far-reaching implications for the spatial and temporal control of gene expression patterns. Additionally, fluctuations in the stability of transcribed mRNA post-splicing can influence the patterns of gene expression during differentiation.^22,23^

In this study, we delve into the intricate regulatory network involving Pol II recruitment, Pol II pausing, chromatin looping, transcription factor binding, and post-transcriptional regulation, shedding light on the sophisticated mechanisms governing regionalized gene expression in the intestinal epithelium. Additionally, we zoom in on the impact of transcription factors such as HNF4 on Pol II recruitment, elucidating its role in bridging the intestinal epigenome with gene expression.

## RESULTS

### Dynamic Pol II recruitment drives differential gene expression during intestinal differentiation

Thousands of genes are differentially expressed in the context of intestinal differentiation, as evidenced by RNA-seq data acquired from isolated crypt and villus epithelium.^13^ Our analysis of this data has identified 1663 transcripts that displayed enrichment in villi, and 2313 transcripts exhibiting enrichment in crypts (log2 fold change [l2FC] > 1 or < -1, FDR < 0.05, FPKM > 1) (Fig. 1A). However, the contribution of RNA polymerase II (Pol II) dynamics during intestinal differentiation has not been systematically explored. Changes in Pol II recruitment, pause-release, or downstream mRNA stability could all contribute to differentiation-specific gene expression. To address this knowledge gap, we conducted chromatin immunoprecipitation sequencing (ChIP-seq) analysis in isolated duodenal villus or crypt cells, using an antibody targeting total RNA polymerase II (Fig. 1B). ChIP replicates from each cell type exhibited a high degree of correlation (Pearson coefficient > 0.9, Fig. S1A). 6429 genes exhibited detectable Pol II signal-over-noise (Fig.S1C-E, Table S1), as determined using size-matched random regions to calculate background signal (Fig. S1B). Individual replicates across the 6429 genes demonstrated a comparable distribution pattern across the gene bodies, wherein enhanced signal was generally observed at transcriptional start sites (TSS). This signal gradually decreased towards the transcriptional end sites (TES), and its strength further declined upon entering intergenic regions (Fig. S1D-F), consistent with other reports of Pol II distribution.^24,25^ Genomic browser tracks of housekeeping genes, *Actb* and *Gapdh*, provided visual validation of the signal quality and robustness inherent in the ChIP experiment (Fig. S1F).

**Fig. 1.**
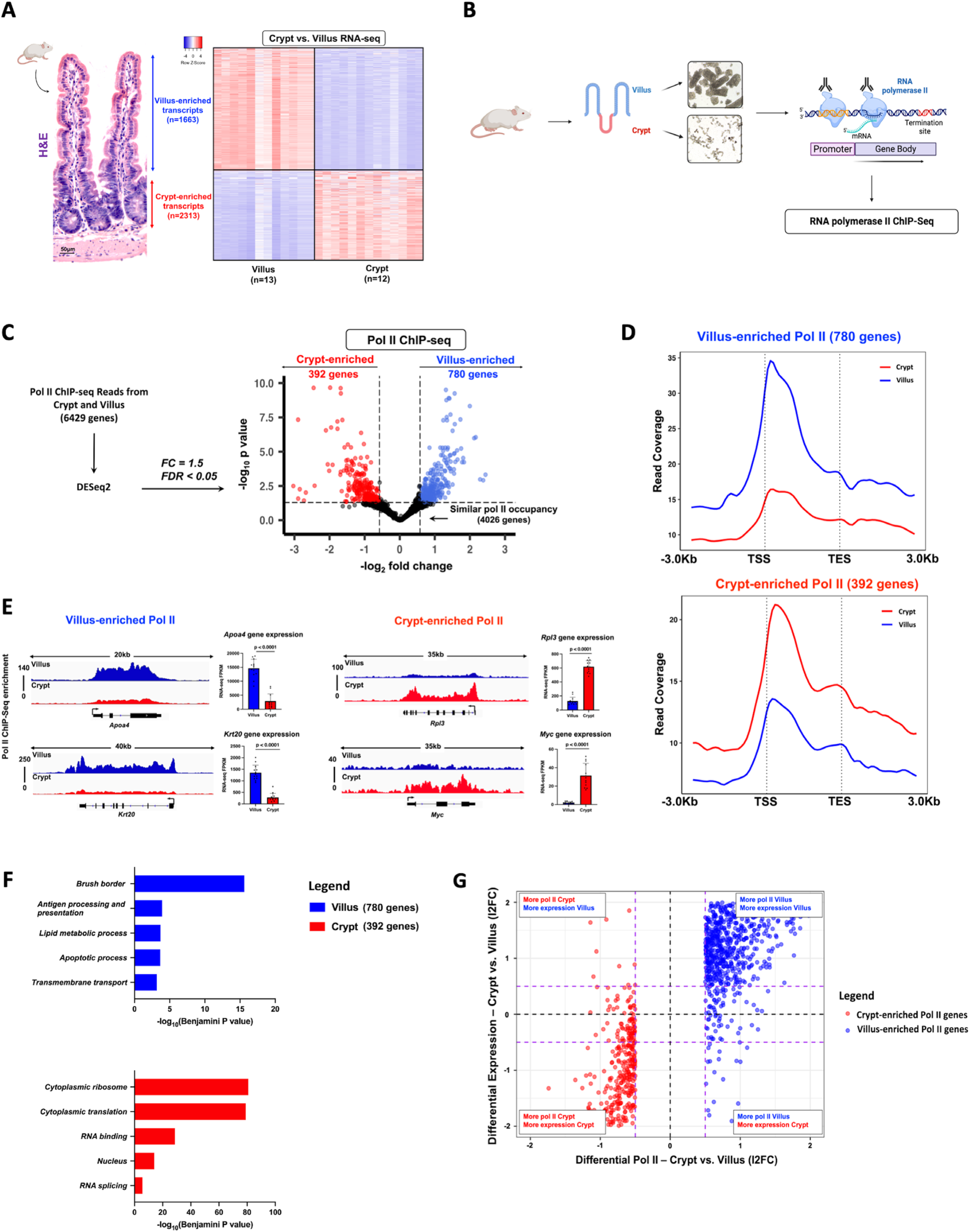
Dynamic Pol II occupancy correlates with dynamic gene expression during intestinal differentiation. **(A)** H&E shows crypt and villus structure from WT mice. Images are representative of 3 biological replicates. Heatmap of villus-enriched and crypt-enriched transcripts reveals dynamic gene expression patterns during differentiation from crypts onto villi (Crypt vs. Villus RNA-seq, n = 13 villi and 12 crypts; DESeq2: l2FC > 1 or < -1, FDR < 0.05, FPKM > 1; GEO: GSE133949). **(B)** Experimental design for Pol II ChIP sequencing from isolated duodenal crypt and villus cells (n=3 biological replicates) **(C)** Volcano plot of differential Pol II occupancy between villus and crypt cells (n = 3 biological replicates). Significant Pol II occupancy was called with DESeq2 (l2FC > 0.58 or < -0.58; FDR < 0.05). Genes with significant Pol II enrichment in the villus and crypt were identified as blue and red points, respectively. Points in black represent genes with similar Pol II binding patterns in both cell types. 780 genes in the villus and 392 genes in the crypt exhibit differential Pol II occupancy (See Table S2). **(D)** Metagene plots show the average signal profiles of Pol II in genes with villus-enriched or crypt-enriched Pol II (differential Pol II; DESeq2, l2FC > 0.58 or < -0.58; FDR < 0.05). **(E)** Functional annotation (DAVID) of villus-enriched Pol II genes and crypt-enriched Pol II genes. p values were calculated using DAVID (See full table in Table S2). **(F)** Examples of differential Pol II binding to gene loci as illustrated using merged Pol II ChIP-seq replicate data. Villus tracks are depicted in blue and crypt tracks are depicted in red. Loci are indicated above, data visualized using IGV. Bar plots show FPKM values for each gene in crypt and villus derived from RNA-seq (GEO: GSE133949). FPKM data is presented as mean ± s.e.m. (n = 13 villi and 12 crypts, two-sided Student’s *t*-test). **(G)** Quadrant plots comparing Pol II-enriched genes with crypt vs. villus differential expression patterns. The red and blue points indicate crypt- and villus-enriched Pol II gene sets respectively (differential Pol II; DESeq2, l2FC > 0.58 or < -0.58; FDR < 0.05), which display large magnitude fold changes in Pol II occupancy and differential expression (crypt vs. villus RNA-seq, DESeq2 l2FC > 1 or < -1, FDR < 0.05, FPKM > 1; GEO: GSE133949). The dotted purple lines show a fold change cut-off of 1.5.

To test whether Pol II is dynamically recruited to different genes during intestinal differentiation, we employed DESeq2 to quantitatively determine differences between villus and crypt as a function of Pol II ChIP read counts per gene. Using an FDR cut-off of 0.05 and at least 1.5-fold difference, we identified two subsets of 780 and 392 genes with significantly enriched Pol II in the villus versus crypt, respectively (Fig. 1C, Tables S2, S3). These gene subsets exhibited greater compartment-specific Pol II occupancy both at the transcription start site (TSS) and across the gene body (Fig. 1D, Fig. S2A). Illustrative examples of differentially enriched Pol II genes include *Apoa4* and *Krt20* in the villus and *Rpl3* and *Myc* in the crypts (Fig.1E). We see in these examples that differential Pol II recruitment and occupancy also corresponds to differential expression of steady-state mRNA as evidenced by the significantly elevated transcripts detected by RNA-seq. Functional annotation of villus-enriched Pol II gene sets demonstrated enrichment of mature enterocyte properties such as brush border and lipid metabolism, whereas genes with enriched Pol II in crypts had functions associated with a proliferative phenotype (Fig. 1F, Table S2). These observations suggest that Pol II recruitment to a gene promoter is highly distinct and is pivotal in controlling cell-specific intestinal functions. To more broadly examine whether dynamic Pol II occupancy correlated with dynamic gene expression during differentiation, we examined changes in Pol II read counts versus changes in corresponding RNA transcripts for genes exhibiting dynamic Pol II recruitment during differentiation. We saw a robust association between dynamic Pol II recruitment and steady-state gene expression (Fig.1G, Fig. S2B), with nearly 92% of villus-enriched Pol II genes, and 85% of crypt-enriched Pol II genes also showing significant mRNA enrichment in their respective compartment (Fig. S2C). These results indicate that dynamic Pol II recruitment is a major driver of the extensive transcriptomic changes that occur during intestinal differentiation.

### A subset of genes exhibits dynamic regulation at the post-transcriptional level

While dynamic Pol II recruitment seemed to explain a large proportion of intestinal gene expression changes, there were 4026 genes which had detectable Pol II ChIP signal but did not make our DESeq2 cut-offs (Fig.1C, Table S3). We categorized this gene set as having similar Pol II occupancies between crypt and villus cells, although we felt most genes in this set still showed differential Pol II levels that did not reach statistical significance (genes indicated by black dots, Fig.1C). Within the set of 4026 genes, a subgroup of 700 genes exhibited differential expression (FPKM > 1, log2 fold change [FC] > 1 or < 1, FDR < 0.05) despite sharing similar Pol II occupancy levels, suggesting alternative mechanisms of transcript regulation at these loci (Fig. 2A-B, Table S2). A metagene analysis on these 700 genes confirmed similar Pol II binding profiles spanning from the TSS to the TES, with nearly overlapping confidence intervals (Fig 2C). Given the uniformity in Pol II recruitment and occupancies across these genes in both crypts and villi, we became intrigued by the possibility that the distinct expression patterns observed in this subgroup might be attributed to post-transcriptional mechanisms, such as alterations in mRNA stability. We examined changes in mRNA stability using the crypt vs. villus RNA-seq data described above. Intronic reads in RNA-seq data are an indicator of newly transcribed, unspliced transcripts, whereas exonic reads should be present in both spliced and unspliced transcripts. We estimated the differences in mRNA stability across the crypt-villus axis using DiffRAC, which is designed to calculate the abundance of *de novo* transcripts versus mature mRNAs using the relative quantities of intronic reads (pre-mRNA) versus exonic reads (mature mRNA).^26^ Analyzing the read distribution across gene exons and introns under distinct conditions enables us to discern whether the transcript remains stable in those conditions (Fig. 2D). For this analysis, we removed 30 genes from the 700 which either had no introns or had zero intron counts in all replicates across both crypts and villi. In these 670 genes, we observed a wider spread in effect size estimates for differential stability than for differential Pol II (Fig. 2E, Table S3). To put this into perspective, nearly 22% of the 670 genes exhibited differential stability at a fold-change of 1.5 compared to none at the same filter for differential Pol II (Fig. 2E). However, a similar analysis with the set of genes showing differential Pol II recruitment (1172 villus- and crypt-enriched Pol II-bound genes) showed a similar fraction (18%) of genes exhibited differential mRNA stability in addition to differential Pol II recruitment (Fig. S3A, Table S3). Thus, while dynamic mRNA stability contributes to differential expression during differentiation, it does not appear unique to genes lacking dynamic Pol II recruitment. We observed that 80% of the villus-enriched transcripts and 72% of the crypt-enriched transcripts within the main set of 670 genes exhibited a compartment-specific rise in mRNA stability (Fig. S3B). Gene Ontology (GO) terms associated with transcripts that are stable and differentially expressed in crypts predominantly pertain to proliferation, whereas those linked with transcripts stable and enriched in villi are primarily related to immunity and lipid metabolism (Fig. S3C, Table S2). To further explore the influence of stability on the expression of genes specific to differentiation, we examined two genes responsible for promoting barrier health and providing protection against colitis (*Muc13*^27,28^ and *Sdc4*^29^) (Fig. 2F-G). *Muc13* and *Sdc4* each exhibit strikingly similar Pol II distributions in both the villus and crypt compartments, despite their mRNA transcript enrichment in only one of these compartments (*Muc13* in villus [Fig. 2F] and *Sdc4* in crypts [Fig. 2G]). Interestingly, there are minimal variations in intronic reads for both genes relative to the changes in exonic reads, indicating similar transcription rates but more substantial changes in the abundance of spliced mRNA. This finding suggests that the changes in crypt-villus expression levels of *Muc13* and *Sdc4* can primarily be attributed to post-transcriptional effects. In summary, our study indicates that mRNA transcript stability serves as a mechanism regulating dynamic expression during intestinal differentiation in a subset of genes.

**Fig. 2.**
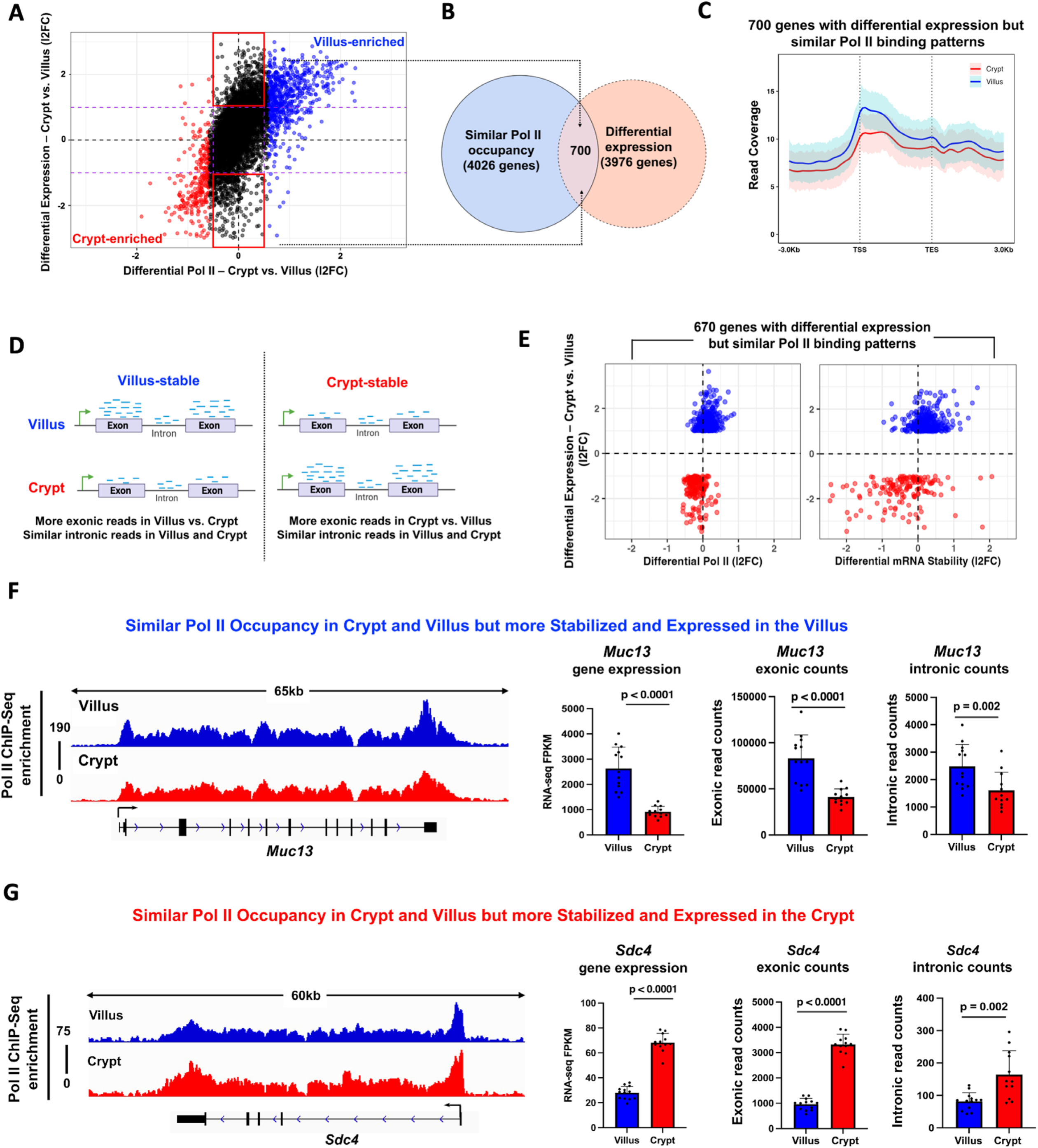
A subset of genes with similar Pol II expression yet differential expression are regulated post-transcriptionally. **(A)** Quadrant plots comparing 6429 genes (from Fig. S1) with gene expression from crypt vs. villus RNA-seq (GEO: GSE133949). The red and blue points indicate crypt- and villus-enriched Pol II gene sets respectively (differential Pol II; DESeq2, l2FC >0.58 or < -0.58; FDR < 0.05). Black points indicate genes which have similar Pol II signal (differential Pol II; DESeq2, l2FC < 0.58 or > - 0.58). Black points highlighted with red boxes indicate genes with similar Pol II occupancy yet are differentially expressed (crypt vs. villus RNA-seq; DESeq2 l2FC > 1 or < -1, FDR < 0.05, FPKM > 1; GEO: GSE133949). The dotted purple lines show a fold change cut-off of 2. **(B)** Venn diagram shows there are 700 genes with similar Pol II occupancy and differential expression between crypt and villus cells (See Table S2). **(C)** Metagene profile shows similarity in Pol II occupancy patterns between crypt and villus in the 700 genes. Confidence bands were generated at 95%. **(D)** Schematic showing principle behind DiffRAC^26^, a computational framework to assess differential mRNA stability based on differential exonic and intronic read counts. **(E)** Plot shows 670 genes (genes without introns were excluded) with similar Pol II occupancy yet differential expression are more likely to be regulated by differential mRNA stability than any differences in Pol II, as evidenced by comparison with differential crypt-villus expression (differential mRNA stability: crypt vs. villus RNA-seq, DiffRAC DESeq2, l2FC > 0 or < 0; differential gene expression: crypt vs. villus RNA-seq, DESeq2 l2FC > 1 or < -1, FDR < 0.05, FPKM > 1; GEO: GSE133949) (See Table S3). **(F)** and **(G)** Representative examples of genes showing similar Pol II occupancies (genomic tracks) yet significantly different exonic counts (bar plots). Villus-enriched is shown in (F) and crypt-enriched is shown in (G). Loci are indicated above, data visualized using IGV. Villus tracks are depicted in blue and crypt tracks are depicted in red. Bar plots depict FPKM values, exonic counts and intronic counts for each gene in crypt and villus derived from RNA-seq (GEO: GSE133949). The data are presented as mean ± s.e.m. (n = 13 villi and 12 crypts, two-sided Student’s *t*-test).

### Dynamic pause-release of Pol II governs regulation of genes during crypt-villus transitions

The dynamic pause-release mechanism of Pol II serves as a rapid tool for finely adjusting gene expression in response to evolving cellular conditions^18,21^. We contemplated the possibility that there might be genes which achieve higher expression levels in the villus by transitioning from a paused and unexpressed state in the crypts. Leveraging Pol II ChIP-seq data, we binned Pol II reads into two zones: the promoter-proximal region (−100bp to +300bp from TSS) and the gene body (+350bp from TSS to TES). This enabled us to compute the Pausing Index (PI)^30–33^ which gauges the ratio of reads in the promoter-proximal region to those in the gene body, normalized by region length (Fig. 3A). The PI provides insights into Pol II pausing at gene promoters. We initially validated the utility of the Pausing Index (PI) as an analytical tool for assessing promoter-proximal pausing of Pol II (Fig. S4). Genes were classified based on their PI in crypt and villus cells. We observed that genes with a PI > 2 displayed a distinct peak in Pol II signal at the TSS, followed by a decline in signal toward the TES. The sharpness of this TSS peak became more pronounced for genes with PI > 3 and PI > 4. Conversely, genes with a PI < 2 lacked a distinct TSS peak and tended to exhibit a higher signal in the gene body, with a pile-up of Pol II towards the TES, likely due to a slowdown of elongation^34,35^ (Fig. S4). We applied DESeq2 to the calculated Pausing Indices and identified 476 genes with more pronounced pausing of Pol II in the crypts (log2 fold change [l2FC] < 0.58, p < 0.05) (Fig. 3B, Tables S2-S3). Of these crypt-paused/villus elongating genes, 280 genes displayed higher mRNA expression in the villus, suggesting that they could be regulated by changes in pause-release between crypts and villi (Fig. 3C, Table S2). A metagene analysis of Pol II distribution across these 280 genes reveals an elevated Pol II signal along the gene body in villus cells compared to the crypts (Fig. 3D). However, it also demonstrates a distinct and prominent peak near the TSS in the crypts, suggesting the presence of Pol II pausing in the crypts, but not in the villi. This observation is consistent with pause-release as the regulatory mechanism governing this subset of villus-enriched mRNAs. Functional annotation shows this set primarily encompassed genes with established villus-specific roles, including functions such as nutrient absorption, lipid transport, immune-related activities, and apoptosis. (Fig. 3E, Table S2). An illustrative example showing a gene from this set, *Dqx1,* provides additional confirmation of the heightened Pol II signal near the promoter-proximal region within the crypts. Moreover, there is a general enhancement in the Pol II signal across the *Dqx1* gene body in the villus. Crucially, there is also clear evidence of significantly higher steady-state gene expression in the villus, as evident by the differences in RNA-seq FPKMs (Fig. 3F).

**Fig. 3.**
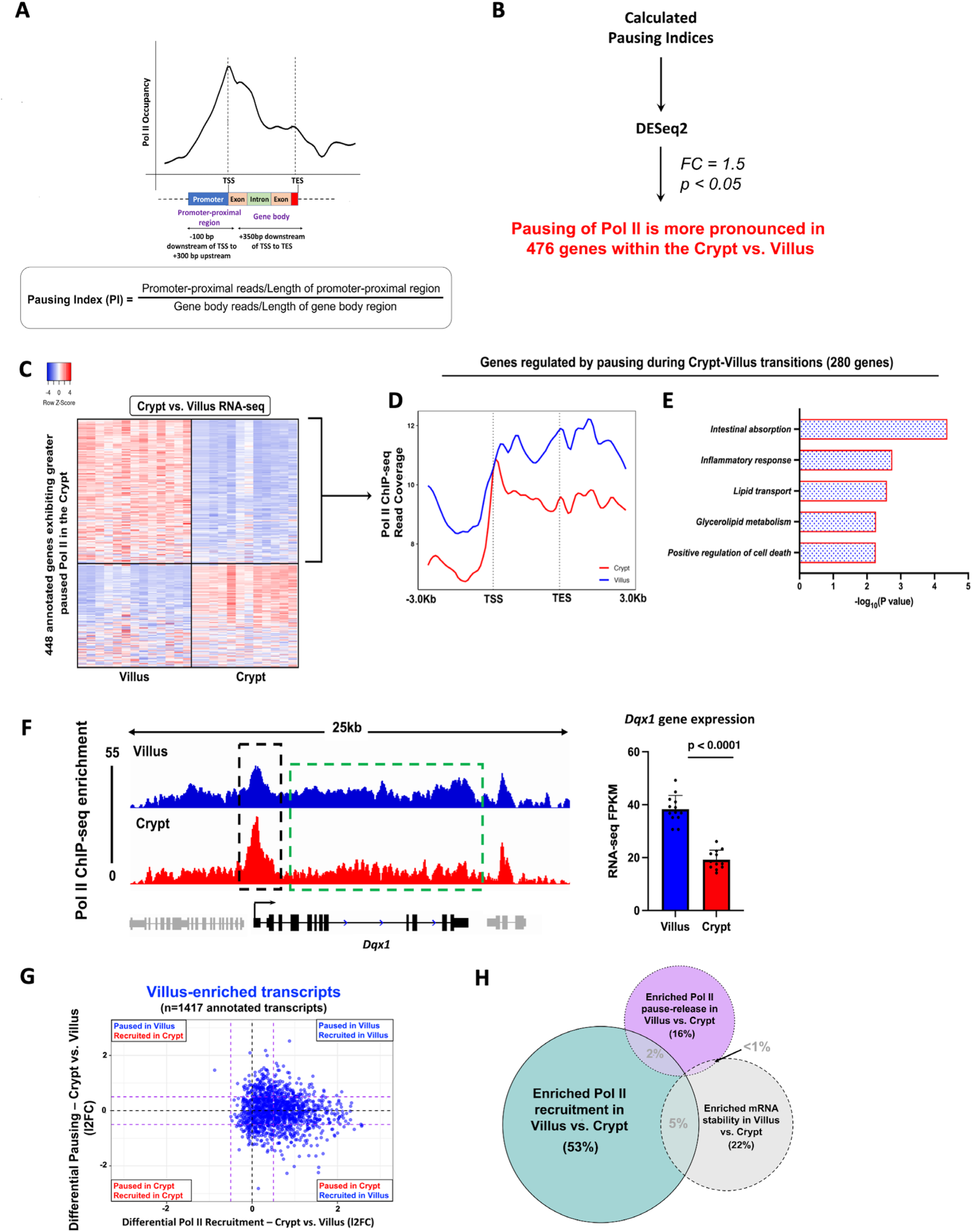
Pause release of Pol II during crypt-villus transitions is an additional gene regulatory mechanism. **(A)** Diagram illustrating the procedure for calculating the Pausing Index (PI) as a measure of promoter-proximal pausing of Pol II. **(B)** Schematic illustrating the approach used to identify 476 genes demonstrating significantly elevated promoter-proximal pausing of Pol II within the crypt (See Table S2). **(C)** Heatmap shows expression patterns of 448 annotated genes which exhibit higher Pol II pausing in the crypt vs. villus (Pol II ChIP-seq; differential PI: DESeq2, l2FC < -0.58, p < 0.05, See Table S2). **(D)** Metagene plot of 280 genes which exhibit greater Pol II pausing in the crypt yet higher expression in the villus, pointing to Pol II pause-release being the major regulatory mechanism. **(E)** Functional annotation of 280 crypt-paused, villus-expressed genes show functional gene classes which preferentially undergo pause release in the villus vs. the crypt (See full table in Table S2). **(F)** An illustration of Pol II occupancy patterns at the *Dqx1* gene. Genomic tracks show elevated Pol II presence at promoters in crypt cells contrasted with heightened Pol II occupancy along gene bodies in villus cells (demonstrated through merged Pol II ChIP-seq replicate data). Villus tracks are depicted in blue and crypt tracks are depicted in red, data visualized using IGV. Bar plots show FPKM values for each gene in crypt and villus derived from crypt vs. villus RNA-seq (GEO: GSE133949). FPKM data is presented as mean ± s.e.m. (n = 13 villi and 12 crypts, two-sided Student’s *t*-test). **(G-H)** Comparison of differential Pol II pausing and differential Pol II occupancy at 1417 genes with villus-enriched transcripts (crypt vs. villus RNA-seq: DESeq2 l2FC > 1, FDR < 0.05, FPKM > 1; GEO: GSE133949) reveals differential Pol II recruitment to be the major gene regulatory mechanism driving differentiation-specific gene expression compared to pause-release or mRNA stability.

In our exploration of differentially expressed genes within the intestinal epithelium, it became apparent that a combination of different regulatory mechanisms was involved in achieving the distinct expression patterns seen during differentiation. To assess the relative impact of various mechanisms contributing to differentiation-induced gene expression, we first plotted the measured differential Pol II recruitment of villus-enriched genes against their corresponding measurement of differential Pol II pausing. We observed that among transcripts enriched in the villus, most of them exhibited increased Pol II recruitment in the villus compared to the crypts. A smaller proportion of villus-enriched transcripts underwent pause-release upon differentiation from crypts onto villi (Fig.3G). 53% of villus-enriched transcripts were primarily governed by differential Pol II recruitment to the promoter (differential recruitment; log2 fold change [l2FC] > 0.58), whereas only 16% were regulated by changes in pausing and pause-release from crypts to villi (differential pausing; log2 fold change [l2FC] < 0.58) (Fig. 3H). Notably, in the villus-enriched transcripts, while increased recruitment of Pol II clearly occurred in the villus, pausing was relatively equally distributed between the crypts and the villus. These observations imply that in addition to genes undergoing pause-release during the crypt-villus transition, other genes might still experience pausing in the villus and engage in pause-release when required. This aligns with previous research suggesting that Pol II pausing is a mechanism that readies genes for prompt initiation of transcriptional elongation.^18,31,36,37^ Finally, 22% of villus-enriched transcripts showed enriched mRNA stability in the villus compared to the crypts (Fig. 3H), providing another mechanism which fine-tunes differentiation-specific gene expression. In summary, it appears that the primary mechanism driving differential gene expression during differentiation across the crypt-villus axis is the differential recruitment of RNA polymerase II to gene promoters.

### Dynamic transcriptional enhancer activities are associated with differential Pol II recruitment during intestinal differentiation

Given that the primary regulatory mechanism in the intestinal epithelium involves the dynamic recruitment of Pol II, we recognized the necessity of incorporating the impact of distal regulatory elements, especially enhancers, to gain a comprehensive grasp of the alterations in gene expression linked to cellular differentiation. We postulated that alterations in Pol II recruitment would be concomitant with shifts in chromatin looping patterns between enhancers and promoters. To explore this, we compared associations between dynamic Pol II recruitment and the frequency of compartment-specific enhancer-promoter looping interactions, as measured by H3K4me3 HiChIP-seq in intestinal villus and crypt cells.^12^ Ligation events captured between distal regulatory elements and each gene promoter were measured in both crypts and villus. Our analysis revealed that 68% of the villus-enriched transcripts (Fig. 4A) and 53.5% of crypt-enriched transcripts (Fig. 4B) displayed a higher frequency of chromatin loops (l2FC > 0.58 or < -0.58) and Pol II occupancy (l2FC > 0.58 or < -0.58) in the corresponding compartment. An illustrative example at the *ApoB* gene promoter shows an increase in villus enhancer-promoter interactions (as measured by H3K4Me3 HiChIP-seq) had a corresponding increase in its Pol II occupancy (as measured by Pol II ChIP-seq) and gene transcript levels (as measured by FPKM values from crypt vs. villus RNA-seq) (Fig. 4C). A similar relationship was observed at a crypt-enriched gene promoter, *Dmbt1* (Fig. 4D). Since there was a correlation between looping events, dynamic Pol II activity, and variations in gene expression, our subsequent focus was on pinpointing the potential transcription factors responsible for driving these dynamic occurrences within the differentiated intestinal epithelium. A total of 5,135 enhancer regions linked with promoters displaying villus-enriched looping events, Pol II occupancy, and gene expression were identified. To discern the transcription factors operating within these 5,135 villus-enriched enhancer regions, we conducted a DNA-binding motif analysis. Notably, the motif corresponding to Hepatocyte Nuclear Factor 4 alpha/gamma (HNF4A/G) emerged as the highest-ranking factor, implicating HNF4 as a key regulator of Pol II dynamics and gene expression in differentiated villi (Fig. 4E, Table S4). Collectively, our findings lead us to the conclusion that the dynamic recruitment of Pol II during intestinal differentiation correlates with dynamic chromatin looping events. This interplay ultimately gives rise to unique gene expression patterns, with this process seemingly under the influence of HNF4 transcription factors.

**Fig. 4.**
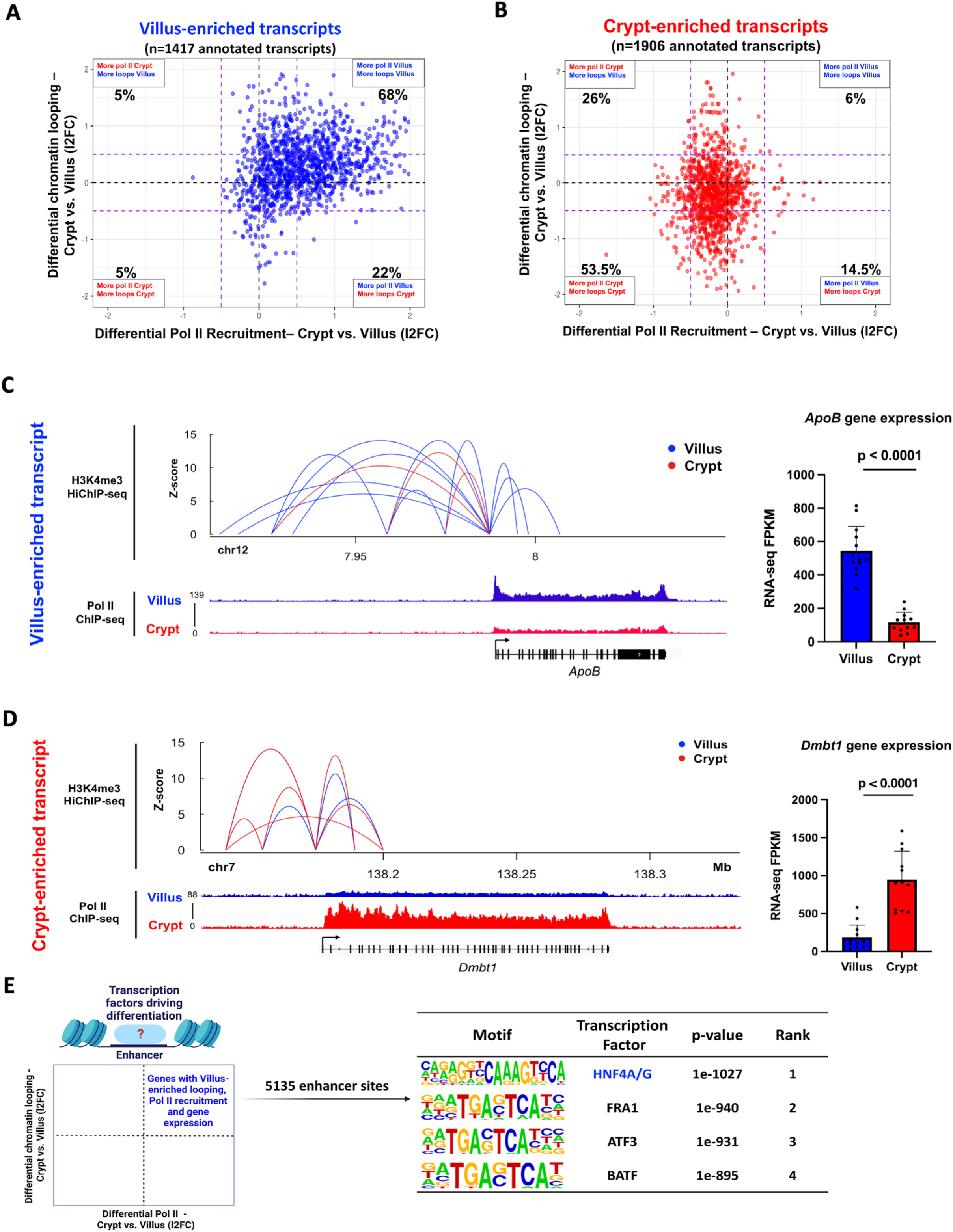
Dynamic transcriptional enhancer activities are associated with differential Pol II recruitment during intestinal differentiation. **(A)** and **(B)** Quadrant plots showing distribution of genes with villus-enriched (A) and crypt-enriched (B) transcripts from RNA-seq data (crypt vs. villus RNA-seq: DESeq2 l2FC > 1 or < -1, FDR < 0.05, FPKM > 1; GEO: GSE133949) with respect to their associated differential enhancer-promoter loops (H3K4Me3 HiChIP-seq: GSE148691) and differential Pol II occupancy (Pol II ChIP-seq). The dotted purple lines show a fold change cut-off of 1.5. **(C)** and **(D)** Gene loci of *ApoB* (C) and *Dmbt1* (D) with corresponding enhancer-promoter loops and Pol II occupancy. Villus tracks are depicted in blue and crypt tracks are depicted in red. All loops shown have q < 0.0001 and counts > 4. Bar plots show FPKM values for each gene in crypt and villus derived from RNA-seq (GEO: GSE133949). FPKM data is presented as mean ± s.e.m. (n = 13 villi and 12 crypts, two-sided Student’s *t*-test). **(E)** Schematic showing the gene set selected for motif analysis using HOMER (genes with higher chromatin looping events, increased Pol II occupancy, and elevated gene expression in the villus) to identify enhancer-bound transcription factors driving differentiation. The highest-scoring motif identified in the analysis corresponds to the transcription factor HNF4. (See full table in Table S4).

### HNF4 transcription factors interact with chromatin remodeling and chromatin looping proteins and are required for Pol II recruitment to differentiation genes

HNF4 transcription factors have roles in activating enhancer chromatin and are essential for the expression of vital villus-enriched genes.^38–40^ Nevertheless, the influence of HNF4 on the dynamics of Pol II remains uncertain, specifically regarding its influence on Pol II recruitment and the subsequent implications for its role as a pro-differentiation factor. To study this, we leveraged a previously generated mouse model which integrates the *Villin-Cre^ERT^*^2^ transgenic allele^41^ with *Hnf4a^flox/flox^* ^42^ on a *Hnf4g^-/-^* background^38^ (hereafter referred to as *Hnf4αγ^DKO)^* (Fig. 5A). Loss of both *Hnf4* paralogs in the intestinal epithelium upon tamoxifen treatment triggered a surplus of proliferative cells, loss of villus integrity and differentiation failure at 4 days after the initial tamoxifen-induced gene knockout (Fig. 5B), consistent with previous reports^38^. We sought to determine the impact of HNF4 loss on the occupancy of Pol II at the genes analyzed above (6429 genes identified in Fig. S1) using ChIP-seq in control and *Hnf4αγ^DKO^* duodenal villus cells at 3 days after tamoxifen treatment, when HNF4 factors are depleted, but the tissue lacks an overt phenotype^38^. Upon loss of *Hnf4* factors, we observed a notable decrease in Pol II recruitment that was particularly pronounced at HNF4-dependent target genes (defined as bound and regulated by HNF4 as measured by HNF4A/G ChIP-seq^38^; within 30 kb of HNF4 binding sites; and differentially regulated in WT vs. *Hnf4αγ^DKO^* RNA-seq^38^) (Fig. 5C, S5A-B, Table S2). For example, within a 210 kb window on chromosome 3, *Pitx2*, a gene not previously identified as being influenced by HNF4, displayed minimal differences in Pol II occupancy between WT and *Hnf4αγ^DKO^* villus cells. In stark contrast, *Enpep*, a known HNF4 target, exhibited a significant reduction in Pol II recruitment upon the absence of HNF4 factors (Fig. 5D).

**Fig. 5.**
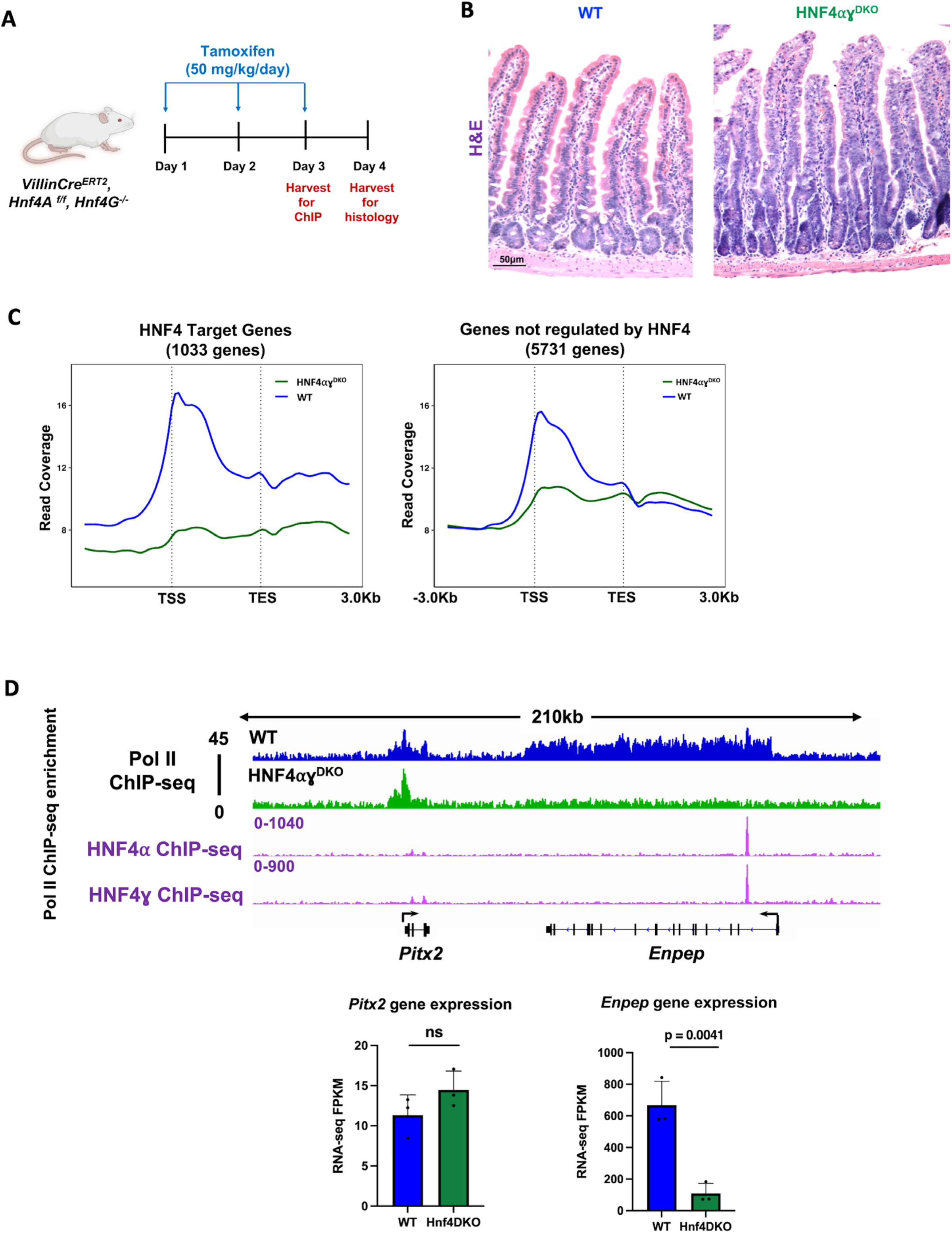
Loss of transcription factor HNF4 compromises Pol II recruitment at its target genes. **(A)** Schematic illustrating the timepoint at which cell collection was performed in *Hnf4αγ^DKO^* mice. For ChIP studies, *Hnf4αγ^DKO^* mice were injected with tamoxifen for 2 consecutive days and harvested the following day. For histology, *Hnf4αγ^DKO^* mice were injected with tamoxifen for 3 consecutive days and harvested the following day. **(B)** H&E staining shows loss of HNF4 paralogs leads to loss of villus integrity and prevalence of elongating crypts (representative of 2 biological replicates). **(C)** Metagene plots show a reduction in Pol II recruitment in *Hnf4αγ^DKO^* villus cells (n = 3). This reduction is particularly pronounced at the genes dependent on HNF4, compared to those which are not (HNF4 targets defined as bound and regulated by HNF4 as measured by HNF4A/G ChIP-seq: within 30 kb of HNF4 binding sites; and differentially regulated in WT vs. *Hnf4αγ^DKO^* RNA-seq; GEO: GSE112946). **(D)** Examples of differential Pol II binding to gene loci in WT and *Hnf4αγ^DKO^* villus within a 210kb window on chromosome 3, as illustrated using merged Pol II ChIP-seq replicate data. WT tracks are depicted in blue and *Hnf4αγ^DKO^* tracks are depicted in green. Loci are indicated above, data visualized using IGV. Bar plots show FPKM values for each gene in WT and *Hnf4αγ^DKO^* cells derived from RNA-seq (GEO: GSE112946). FPKM data is presented as mean ± s.e.m. (n = 3 biological replicates per group, two-sided Student’s *t*-test).

Building on these findings, we sought to identify all genes exhibiting distinct Pol II patterns in wild-type (WT) mice compared to *Hnf4αγ^DKO^* mice. By applying DESeq2 on Pol II ChIP-seq read counts (log2 fold change [l2FC] > 0.58 or < -0.58, FDR < 0.05), we identified nearly 2000 genes which had differential Pol II occupancy between control and *Hnf4αγ^DKO^* villus epithelia (997 WT-enriched Pol II genes; 882 *Hnf4αγ^DKO^* -enriched Pol II genes) (Fig.6A, Tables S2-S3). Clear differential Pol II signals at both genes gaining and losing Pol II ChIP signal were demonstrated in the heatmap (Fig. 6B).

**Fig. 6.**
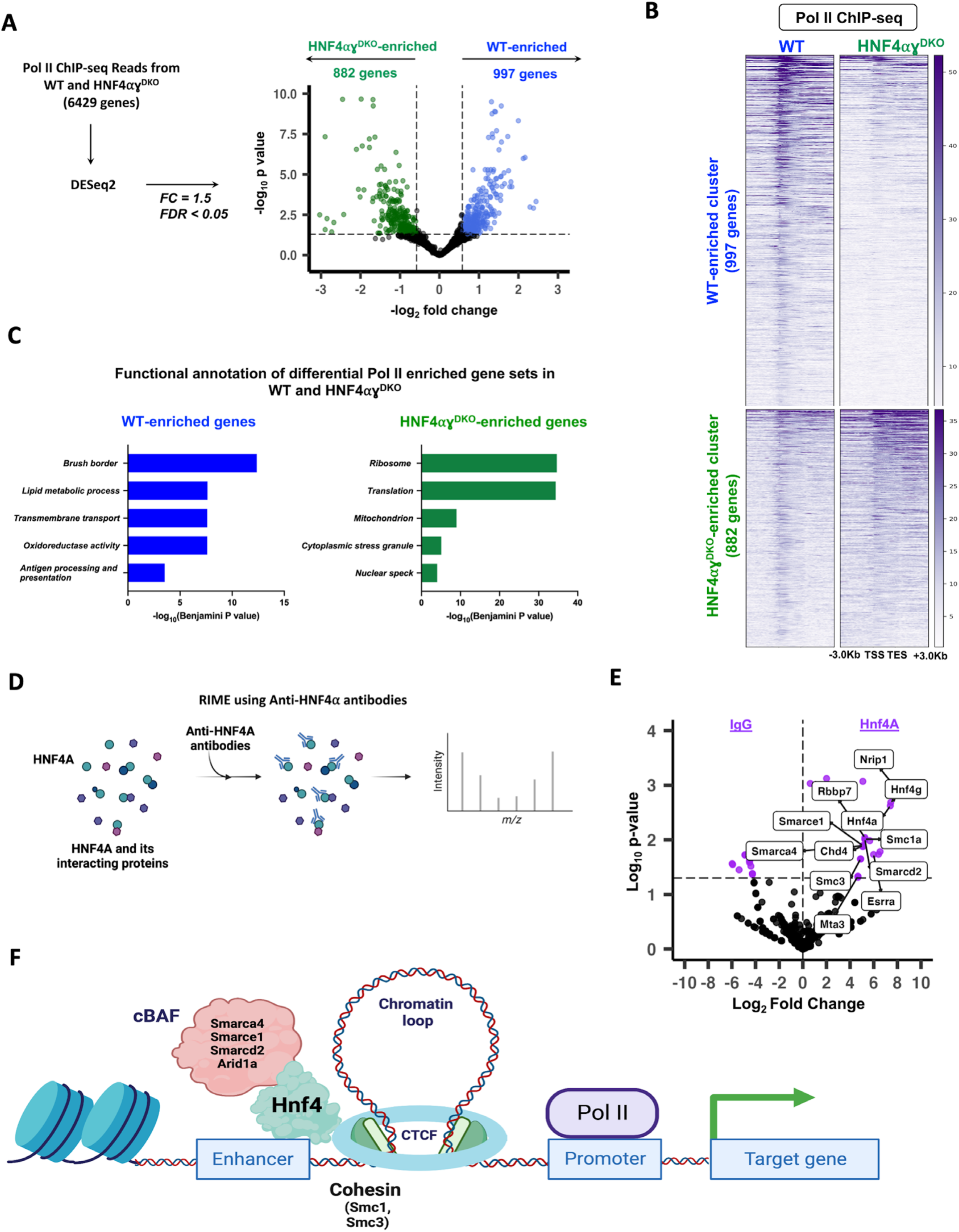
HNF4 fosters open chromatin and enhances Pol II recruitment at genes specific to differentiation. **(A)** Volcano plot of differential Pol II occupancy between WT and *Hnf4αγ^DKO^* villus cells (n = 3 biological replicates). Significant Pol II occupancy was called with DESeq2 (Pol II ChIP-seq: l2FC > 0.58 or < -0.58, FDR < 0.05). Genes with significant Pol II enrichment in WT and *Hnf4αγ^DKO^* were identified as blue and green points, respectively. Points in black represent genes with similar Pol II binding patterns in both cell types. 997 genes in the WT and 882 genes in the *Hnf4αγ^DKO^* exhibit differential Pol II occupancy (see Table S2). **(B)** Heatmap shows the distribution of Pol II signal across the gene loci for 997 and 882 WT- and *Hnf4αγ^DKO^* - enriched Pol II gene sets, respectively (Pol II ChIP-seq: DESeq2, l2FC > 0.58 or < -0.58; FDR < 0.05). **(C)** Functional annotation of differential Pol II gene sets show an enrichment in HNF4-dependent, villus specific functions in WT and a reversion to a crypt-like state in the *Hnf4αγ^DKO^*. p values were calculated using DAVID (see full table in Table S2). **(D)** Schematic for Rapid Immunoprecipitation Mass Spectrometry of Endogenous proteins (RIME) using anti-HNF4⍺ antibodies. **(E)** Differential analysis of peptide counts from RIME directed against HNF4⍺ and IgG in primary mouse epithelium shows that cohesin subunits and peptides of the SWI/SNF complex co-IP with anti-HNF4⍺. Peptide fragments with > 1 spectral count and > 2 unique peptide fragments were used for analysis (DESeq2 of peptide counts, FDR < 0.05) (see Table S5). **(F)** Proposed model suggesting a mechanism by which HNF4 facilitates enhancer-promoter looping and chromatin remodeling, thus providing a conducive environment for Pol II recruitment to a promoter.

Moreover, at the individual gene level, the contrasting Pol II occupancy is clearly visible in the representative examples (Fig. S6A). Functional annotation of genes showing loss of Pol II recruitment in the *Hnf4αγ^DKO^* reveals functions associated with differentiation, whereas those genes gaining Pol II in the *Hnf4αγ^DKO^* are associated with functions in cellular proliferation and stress responses (Fig. 6C, Table S2). We observe a strong correlation between dynamic Pol II occupancy and gene expression changes in the *Hnf4αγ^DKO^* (Fig. S6B-C). We also discovered that approximately 40% of the genes losing Pol II recruitment in the *Hnf4αγ^DKO^* were also bound by HNF4 in ChIP-seq and show decreased RNA levels in the *Hnf4αγ^DKO^*, underscoring the critical role of HNF4 in regulating Pol II recruitment and transcriptional activity during intestinal differentiation (Fig. S6D).

To gain insights into the mechanisms underlying HNF4-mediated Pol II recruitment, we performed RIME (Rapid Immunoprecipitation Mass Spectrometry of Endogenous proteins)^43^ using anti-HNF4α antibodies in primary mouse epithelium (Fig. 6D, Table S5). While analysis of the HNF4 interactome has been performed in cell lines^44,45^ and murine colon^46^, we provide here the first analysis in the differentiated small intestine. Differential analysis of raw peptide counts from IgG and HNF4α co-immunoprecipitated samples led to the identification of 29 proteins which exhibit significant interaction with HNF4α (l2FC > 0.58, FDR < 0.05) (Table S5). Within the top 20 hits we obtained, it is noteworthy that 40% of them were transcriptional regulators associated with the SWI/SNF chromatin remodeling complexes (Fig. 6E, Table S5).

Notably, we also identified interactions with SMC1A and SMC3, integral components of the cohesin protein complex, which plays a critical role in facilitating DNA looping^47,48^ (Fig. 6E). Based on these findings, we propose a model that suggests the following mechanism: HNF4 binding to an enhancer region initiates chromatin remodeling facilitated by the SWI/SNF complex. This remodeling leads to the formation of a chromatin loop between the enhancer and gene promoter, thus, creating a favorable environment for the recruitment of Pol II to a target gene regulated by HNF4 (Fig. 6F). By influencing chromatin accessibility, interacting with transcriptional regulators, and facilitating the assembly of transcriptional complexes, HNF4 ensures proper Pol II recruitment and subsequent gene expression driving intestinal differentiation.

## DISCUSSION

The regulation of gene expression during tissue differentiation is a multifaceted process, and the relative significance of Pol II recruitment, pause release, and mRNA stability can vary across different tissue types and developmental contexts. While *de novo* Pol II recruitment is a fundamental step in transcription and often contributes to differential gene expression during differentiation, other mechanisms may take precedence in different tissues. For instance, in embryonic stem cells, the dynamics of Pol II pause release and elongation rates are prominent as cells differentiate into various lineages.^37^ In contrast, during skeletal muscle regeneration, both Pol II pause-release and mRNA stability have been implicated in sculpting the muscle-specific transcriptome.^49,50^ In immune cells, such as T cells, mRNA stability and post-transcriptional regulation are prominent mechanisms for orchestrating rapid and precise gene expression changes in response to immune challenges.^51^ These diverse examples underscore the tissue-specific and context-dependent nature of gene expression regulation for maintaining cellular homeostasis.

The intestinal epithelium, characterized by its constant renewal, presents a valuable yet underexplored model system for studying the mechanisms of transcriptional differentiation. Maintaining the equilibrium between epithelial cell proliferation and differentiation constitutes a critical physiological response during viral infections^52^, periods of fasting^53^, or during the consumption of fat-rich diets^54^. Deviations from the normal levels of differentiation can render individuals more susceptible to diseases pathologies, including metabolic syndrome^54^ and cancer.^54–56^ Understanding the mechanisms governing differentiation is also of paramount importance in the field of regenerative medicine, where the delicate balance between cell expansion and the generation of fully functional, differentiated tissue is pivotal to success. The intestinal crypt-villus unit plays a vital role in upholding intestinal equilibrium and overall health.

Nonetheless, the precise gene regulatory mechanisms during the transition from crypts onto villi remain unclear. Our research has extensively examined the impact of diverse Pol II patterns driving alterations in gene expression and cell-specific functions along this axis. A key result from our study was that the driver of gene expression changes across the crypt-villus axis stemmed from the dynamic recruitment of Pol II to gene promoters. Additionally, smaller subsets of genes were regulated through changes in Pol II pausing and post-transcriptional mechanisms. Moreover, we demonstrated that the dynamic interplay of chromatin looping also contributes to the recruitment of Pol II to intestinal gene promoters. This phenomenon is substantially influenced by the transcription factor HNF4 in differentiated villus cells.

Current research in intestinal biology frequently highlights the BMP signaling pathway as the central force behind intestinal differentiation^57^. This assertion is grounded in sophisticated studies employing transgenic expression of BMP antagonists leading to ectopic crypt formation in the villus^58,59^. A focal point for future investigations could include the molecular mechanisms of how the BMP signaling pathway activates differentiation-specific gene expression. SMAD4 acts as the transcriptional effector of BMP and mutations within it are among the most prevalent in colon cancers.^56,60–63^ SMAD4 and HNF4 mutually activate each other’s expression and subsequently bind to and activate genes associated with enterocyte differentiation.^38^ Yet, the impact of this interaction on *cis*- and *trans*-regulatory factors within the differentiated intestinal genome remains undocumented. It raises intriguing questions about whether HNF4’s role in mediating open chromatin might facilitate the access of other pro-differentiation factors like SMAD4, to DNA, thereby modulating the expression of differentiation genes. Or if they collaborate in processes such as chromatin opening, recruitment of Pol II or the release of transcriptional pauses. These findings have the potential to uncover crucial transcriptional mechanisms of pro-differentiation factors in normal intestinal physiology, shedding light on their relevance to conditions such as Inflammatory Bowel Disease (IBD) and cancer.

In conclusion, this comprehensive understanding of the regulatory network underlying dynamic crypt-villus gene expression patterns not only advances our knowledge of intestinal biology but also provides insights into the broader mechanisms of tissue specialization and homeostasis. Further investigations into intestinal regulatory mechanisms promise to uncover additional layers of complexity and deepen our understanding of the remarkable adaptability and functionality of the intestinal epithelium.

## ACKNOWLEDGEMENTS

This research was funded by grants from the National Institutes of Health (NIH) to M.P.V. (R01DK121915 and R01DK126446); K.V. was supported by an American Heart Association pre-doctoral fellowship (906006). S.K. was supported by a Rutgers DLS Summer Undergraduate Research Fellowship. L.C. was supported by grants from Start-up Research Fund of Southeast University (RF1028623015), National Natural Science Foundation of China (32270830) and Jiangsu Provincial Science Fund for Distinguished Young Scholars (BK20230026). Figures 1A, 1B, 2D, 4E, 5A, 6D, 6F, and S1C were created with the aid of Biorender.com. The authors acknowledge the Office of Advanced Research Computing (OARC) at Rutgers University for providing access to the Amarel cluster and associated research computing resources which have contributed to the results reported here.

## AUTHOR CONTRIBUTIONS

K.V. designed the study, performed benchwork, sequencing data processing, and bioinformatics; collected and analyzed the data; and wrote the manuscript. S.K. contributed to sequencing data processing, bioinformatics, and data analysis. L.C. performed benchwork and contributed to RIME data processing. M.P.V. conceived, designed, and supervised the study; contributed to sequencing data processing and data analysis, and wrote the manuscript.

## METHODS

### Mice

*Villin-Cre^ERT^*^2^ transgene^41^, *Hnf4α^f/f^* ^42^ and *Hnf4γ^Crispr/Crispr^* ^38^ alleles were integrated to generate conditional compound mutants and controls. Experimental *Villin-Cre^ERT^*^2^ *Hnf4* mutant mice (8–12 weeks old) were intraperitoneally administered tamoxifen at 50 mg/kg/day (Sigma, T5648). Wild-type, cage mate controls were administered corn oil vehicle (Sigma, C8267) at the same dosage. For ChIP-seq experiments, duodenal tissue was collected after 2 consecutive days of tamoxifen or vehicle treatment. For histology experiments, tissue was collected after 3 consecutive days of tamoxifen or vehicle treatment. For RIME, villi from the proximal half of the small intestine from C57BL/6 mice were collected. Mice of both sexes were used in all experiments. Mouse protocols and experiments were approved by the Rutgers Institutional Animal Care and Use Committee and all relevant ethical regulations were followed. All samples were collected between 9:00am to 12:00pm to avoid circadian variability.

### Epithelial Cell Isolation

The initial third of the small intestine (duodenum) was dissected and opened longitudinally to wash the inter-lumen space and expose epithelial cells. Tissue was cut into 1-inch pieces and rinsed with ice-cold PBS. Tissue was treated with 3mM EDTA/PBS for 25 minutes for mutant mice and 35 minutes for wild-type, with changes in EDTA solution at the initial 5- and 10-minute time points. Mechanical force was applied to release the epithelial cell layer from the underlying mesenchyme. Crypts and villi were separated with a 70uM filter. The smaller crypts pass through the pores of the filter while the larger villi stay on top. Both crypts and villi were collected from wild-type mice, while only villi were collected from *Hnf4* mutant mice. Cells were washed twice with ice-cold PBS and pelleted by centrifugation at 300 rcf at 4°C.

### Histology and immunostaining

Intestinal tissues were fixed overnight in 4% paraformaldehyde at 4°C, washed with PBS, and dehydrated through ascending alcohols before paraffin embedding. Five-micrometer paraffin sections were used for Hematoxylin & Eosin staining (H&E). The staining was imaged with a Lumenera INFINITY3 camera and Infinity Analyze imaging software (v6.5.6).

### Pol II ChIP-seq

Crypt and villus cell pellets were cross-linked with 1% formaldehyde (Sigma, F8775) at 4°C for 10 min and then at 25°C for 50 min. Cells were pelleted and washed with ice-cold PBS twice by centrifugation at 300 rcf at 4°C. Fixed cells were lysed in 3X volume of lysis buffer (1% SDS, 10mM EDTA, 50mM Tris-HCl pH 8.0, 1X protease inhibitor cocktail (G-Biosciences, 786-433). Cells were sonicated for 35 minutes using a Diagenode Bioruptor to shear chromatin to 200-500bp. Protein A/G beads (Invitrogen, 10001D and 10004D) were washed with 1% BSA in PBS and pre-loaded with Anti-RNA polymerase II antibody (Millipore,05-623, Lot 3506610). Antibody-conjugated beads were washed twice with 1% BSA and incubated with sheared chromatin in dilution buffer (1% Triton X -100, 2mM EDTA, 150mM NaCl, 20mM Tris-HCl, pH 8.0). The final concentration of SDS in the sonicate was 0.26%. Chromatin and beads were incubated on a rotator at 4°C overnight. Chromatin-bound, antibody-conjugated beads were collected by magnetic separation. Unbound chromatin was discarded. Beads were washed five times with RIPA buffer (50mM HEPES pH 7.6, 1mM EDTA, 0.7% Sodium deoxycholate, 1% NP-40, 0.5M LiCl), and once with TE buffer (10mM Tris, 0.1mM EDTA). Cross-links were reversed in buffer containing 0.1M NaHCO3 and 1% SDS overnight at 37°C to release ChIP DNA. DNA was column purified using the MinElute PCR purification kit (Qiagen, 28004) and quantified using Picogreen (Invitrogen, P7581). Libraries were prepared using the ThruPLEX DNA-Seq kit (Takara, R400675) and DNA Unique Dual Index Kit (Takara, R400666). Paired-end sequencing of ChIP-seq libraries was performed (150 bp) to a depth of at least 30 million reads on the Illumina NovaSeq 6000 platform, demultiplexed and converted to fastq format.

### ChIP-seq Data Processing

FastQC (v0.11.3)^64^ was used to check the quality of raw sequencing reads. Alignment to the mouse reference genome (MGSCv37/mm9) was done using bowtie2 (v2.2.6)^65^ and alignments were sorted using Samtools (v0.1.19)^66^. When required, Picard’s MergeSamFiles was used to merge replicate bam files (http://broadinstitute.github.io/picard). deepTools2 bamCoverage (v2.4.2)^67^ was used to generate RPKM-normalized bigwig files from individual and merged bam files. The Integrative Genomic Viewer (v2.8.13)^68^ was used to visualize normalized bigwig tracks. The BEDTools utility (v2.17.0)^69^ was used to merge, intersect, or subtract the intervals of bed files. Enriched ontologies were identified with DAVID (v2021).^70^ ^71^ For generation of heatmaps and metagene profiles, bigwigs were first quantile-normalized using Haystack (v0.4.0).^72^ deepTools2 (v2.4.2) computeMatrix, plotHeatmap and plotProfile were used to generate plots at defined genomic regions. multiBamSummary and plotCorrelation from deepTools2 (v2.4.2) were used to compute Pearson correlations showing similarity between replicates.

### RNA-seq and HiChIP-seq Data Processing

RNA-seq data (GSE133949) was re-analyzed using Kallisto (v0.44.0)^73^ and DESeq2 (v1.36.0)^74^ for the R programming language (R v4.2/ RStudio v2022.02.3/ Bioconductor v3.15). GSEA (v4.2.2)^75^ was done on pre-ranked gene lists. Genes with FPKM > 1 were used for further analysis. Heatmapper^76^ and Morpheus (https://software.broadinstitute.org/morpheus) were used for k-means clustering and/or visualization of normalized transcript levels of genes of interest.

For identification of differential loops from H3K4me3 HiChIP-seq data, we considered all loops with q ≤ 0.0001 in at least one replicate and raw counts ≥ 4 in both replicates. DESeq2 (v1.36.0) was applied to identify differential looping events using sequencing counts. Promoters involved in these differential looping events were extracted and annotated using BEDTools (v2.17.0) by intersection with a whole genome promoter bed file (UCSC transcription start site ± 2 kb). The number of loops associated with each gene promoter was summed and averaged between replicates, and DESeq2 (v1.36.0) was run again to identify differential looping between each gene promoter. The R package, Sushi (v1.20.0)^77^ was used to visualize HiChIP-seq looping data. For simplicity, we combined biological replicates for visualizing loops in Sushi.

## Pol II ChIP-seq Bioinformatics Data Analysis

For further analyses, each gene was partitioned into the promoter-proximal region and the gene body. Raw gene-level counts data for each region was obtained using featureCounts from the Subread package (v2.03)^78^. Modified whole-genome annotation files specific to the promoter-proximal region and gene body regions were generated as input for featureCounts. Custom annotation files were generated as follows: 1) From the mm9 transcript-level annotation (RefSeq), TSS coordinates were extracted from forward and reverse strands for all isoforms associated with a gene; 2) SAF annotation format files were generated with only promoter-proximal region (TSS, -100 to +350) and gene body regions (+350 to 500bp before TES); 3) Finally, transcripts shorter than 1000bp were removed. The isoform with the strongest signal at the promoter proximal region was retained for further analyses. For analysis of Pol II recruitment and pausing, transcripts from non-standard chromosomes, non-coding transcripts, and those with less than 18 counts/kb gene length were excluded. For the remaining transcripts, promoter proximal and gene body counts were normalized by region length. Pausing Index (PI) was calculated as the ratio of normalized reads at the promoter proximal region to the normalized gene body reads for each gene.

Differential expression analyses were performed using the DESeq2 package (v1.36.0). For analysis of differential Pol II recruitment, total reads mapped to the full gene were used. For differential pausing analyses, the PI multiplied by a factor of 1000 was used as DESeq2 input.

Data visualization was performed using GraphPad Prism (v9.50) and the R packages ggplot2 (v3.3.6)^79^, ggpubr (v0.6.0)^80^, metagene (v2.8.1)^81^ and eulerr (v7.0.0).^82^

### Differential mRNA stability analysis

Differential mRNA stability analysis was performed as previously described.^26^ Briefly, crypt vs. villus RNA-seq data was aligned to Ensembl GRCm39 version 109 using HISAT2 (v 2.2.0)^83^. Using the CRIES workflow (https://github.com/csglab/CRIES)^84^, reads were mapped to constitutive exonic and intronic regions for transcripts supported by both Ensembl and Havana consortia and quantified with featureCounts (v2.03).

Genes without introns or those with zero intron counts in all replicates across all conditions were removed from subsequent analyses. DiffRAC was used to infer changes in mRNA stability as described previously.^26^

### Transcription factor motif analysis

Enhancers associated with villus-enriched gene sets were identified by employing BEDtools intersect and pairToBed to overlay their promoters with H3K4me3 HiChIP-seq regions. Enhancer locations were further refined by use of a subsequent BEDtools intersect step with villus ATAC-seq peaks^13^ to center enhancer regions within areas of open chromatin. HOMER findMotifsGenome.pl (v.4.8.3) was used to call transcription factor motifs enriched at enhancers.^85^

### Rapid Immunoprecipitation Mass Spectrometry of Endogenous Proteins (RIME)

Villus cells from the proximal small intestine of C57BL/6 mice were collected in ice-cold PBS by scraping the everted tissue with a cover slip. Care was taken to avoid collecting the muscle layer. Cell suspensions were pooled, snap-frozen in liquid nitrogen and submitted to Active Motif for RIME analysis according to published protocols^43^. RIME was carried out using an antibody against HNF4α (Santa Cruz Biotechnology, sc-6556 X, Lot B1015) to identify peptides which interact with HNF4α using mass spectrometry. An isotype-matched IgG antibody was used as control. Analysis was performed in duplicate for each immunoprecipitation. Bioinformatic analysis was performed with the software Scaffold (v5.2.2)^86^ and enriched peptide fragments which showed 2-fold enrichment over IgG controls, > 1 spectral count and > 2 unique peptide fragments were listed as interacting partners of HNF4α.

### Statistical analysis

Data is presented as mean ± s.e.m., and statistical comparisons were performed using two-way analysis of variance (ANOVA) or two-sided Student’s *t*-test at p < 0.05 with GraphPad Prism (v.9.50) or ggplot2 (v3.3.6). Mann–Whitney *U*-test and Kruskal–Wallis test (followed by post hoc Dunn’s test) were used as part of RNA-seq analysis. Bioinformatics-related statistical analysis was performed with the embedded statistics in each package, including DESeq2^74^, GSEA^75,87^, and DAVID^70,71^.

### Data and Code Availability

ChIP-seq data from this publication have been deposited to GEO accession number GSE244918. Mass spectrometry data from the RIME analysis have been deposited into the MassIVE database with the dataset identifier MSV000093071 (ftp://MSV000093071@massive.ucsd.edu). The following datasets from GEO were reanalyzed with our sequencing data: GSE133949^13^ was used to perform RNA-seq analysis of villus-enriched genes and crypt-enriched genes; GSE148691^12^ was used to analyze enhancer-promoter looping across the crypt-villus axis; GSE112946^38^ was used to identify HNF4 binding patterns in intestinal epithelial cells. All newly generated code used in this analysis as well as data files and additional documentation can be found in our GitHub repository at https://github.com/VerziLab/Intestinal-RNA-Pol-II-Dynamics.

**Fig. S1.**
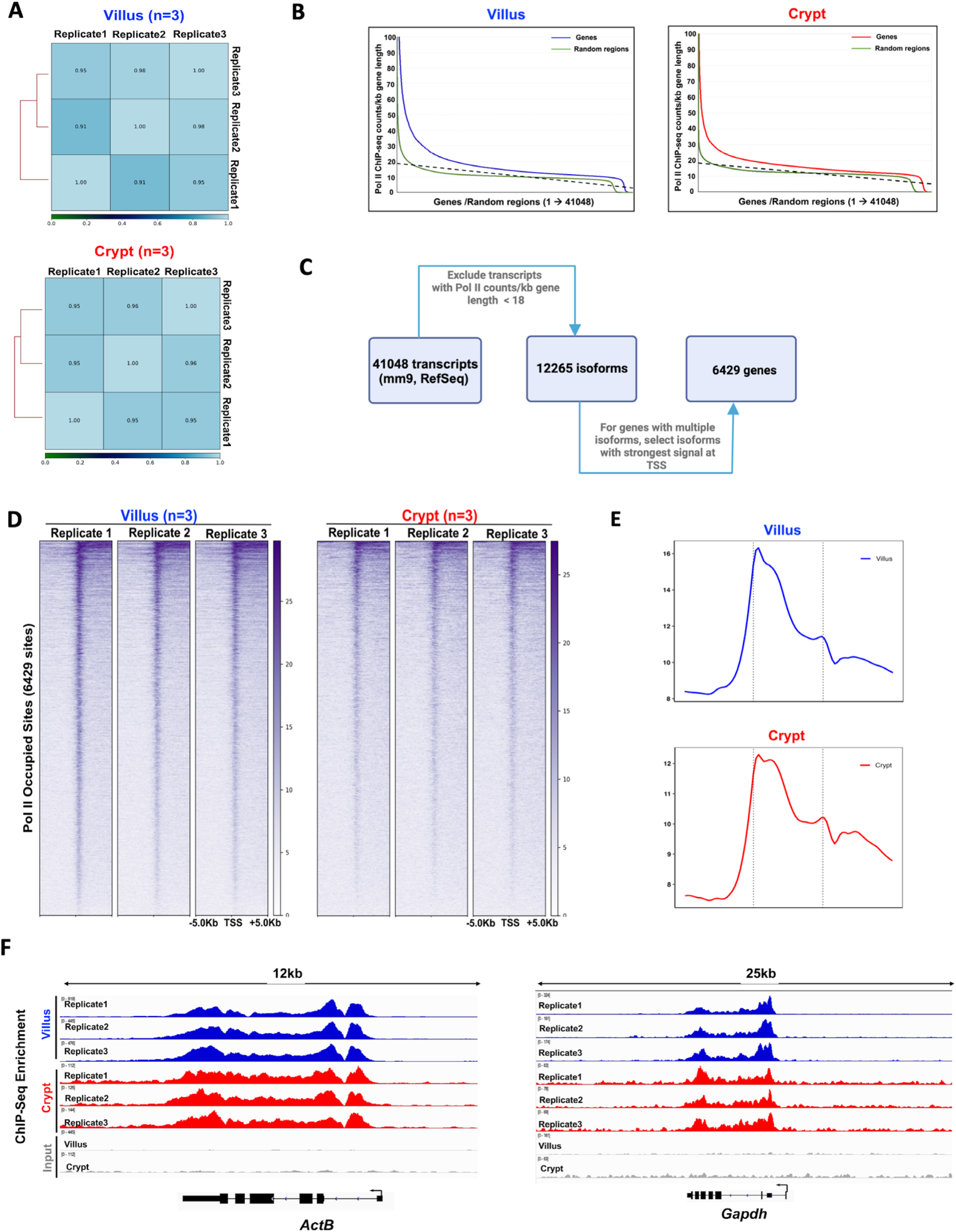
Quality assessment of Pol II ChIP-seq in villus and crypt cells. Related to Fig. 1. **(A)** Correlation plots demonstrate a high degree of similarity between biological replicates in the Pol II ChIP-seq experiment for both villus and crypt cells, with Pearson correlation coefficients exceeding 0.9 in all instances. (B) Comparison of Pol II signal-over-noise in villus and crypt cells. Pol II signal at 41,048 isoforms from the NCBI RefSeq database (Genome build: mm9) versus size-matched random regions were used to calculate background signal. Dotted black line depicts the y-intercept, below which was designated as background and above which was designated as true Pol II signal. Data representative of 3 biological replicates. **(C)** Schematic depicting the gene set selection process for the study, involving the removal of genes displaying background signals and the choice of the isoform with the most robust signal at the TSS. 6429 genes were selected based on a cut-off of > 18 counts/kb gene length (see Table S1). **(D)** Heatmaps show detectable Pol II signal at TSS ± 5kb in all biological replicates of crypt and villus cells across 6429 genes (n=3 biological replicates). **(E)** Metagene profiles show detectable Pol II signal across the gene in biological replicates of crypt and villus cells of 6429 genes (n=3 biological replicates). **(F)** Instances of analogous Pol II binding patterns observed at the gene loci of housekeeping genes, *Actb* and *Gapdh*, across each replicate in both crypt and villus cells. Villus tracks are depicted in blue and crypt tracks are depicted in red. Data was visualized using IGV.

**Fig. S2.**
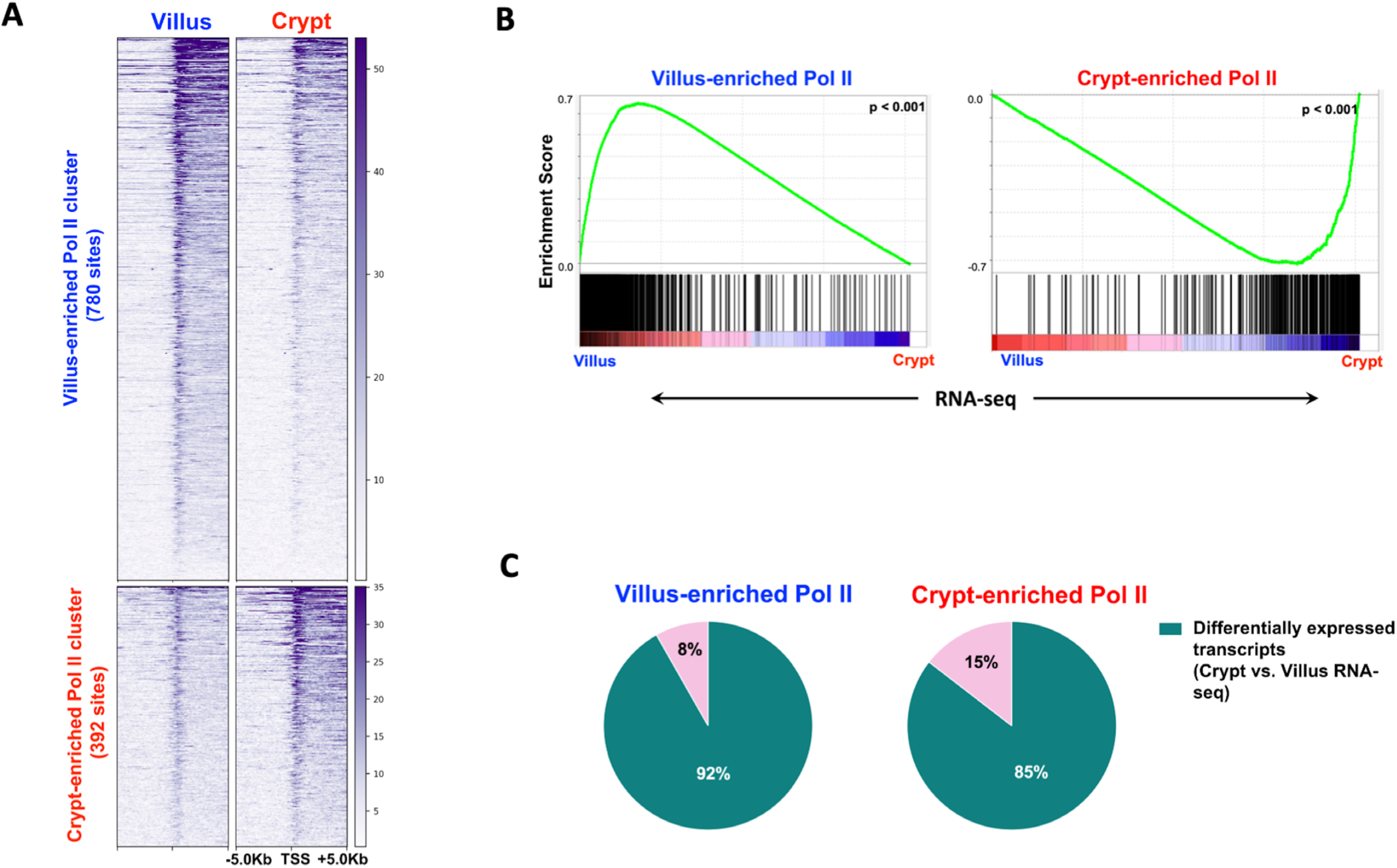
Further characterization of villus-enriched and crypt-enriched Pol II gene sets. Related to Fig. 1. **(A)** Heatmap shows the distribution of Pol II signal at TSS ± 5kb for the 780 and 392 villus and crypt-enriched Pol II gene sets, respectively (Pol II ChIP-seq: DESeq2, l2FC > 0.58 or < -0.58; FDR < 0.05). **(B)** Gene set enrichment analysis (GSEA) reveals that villus-enriched Pol II genes are highly expressed in villus cells, whereas crypt-enriched Pol II genes are highly expressed in crypt cells (Kolmogorov-Smirnov test, p < 0.001). **(C)** Proportion of Pol II enriched genes in villus and crypt which show more steady-state gene expression in the same cell type. In villus cells, 92% of Pol II-enriched genes in the villus are more highly expressed, while in crypts, 85% of Pol II-enriched genes in the crypt display increased expression (crypt vs. villus RNA-seq: DESeq2, l2FC > 1 or < -1, FDR < 0.05, FPKM > 1; GEO: GSE133949).

**Fig. S3.**
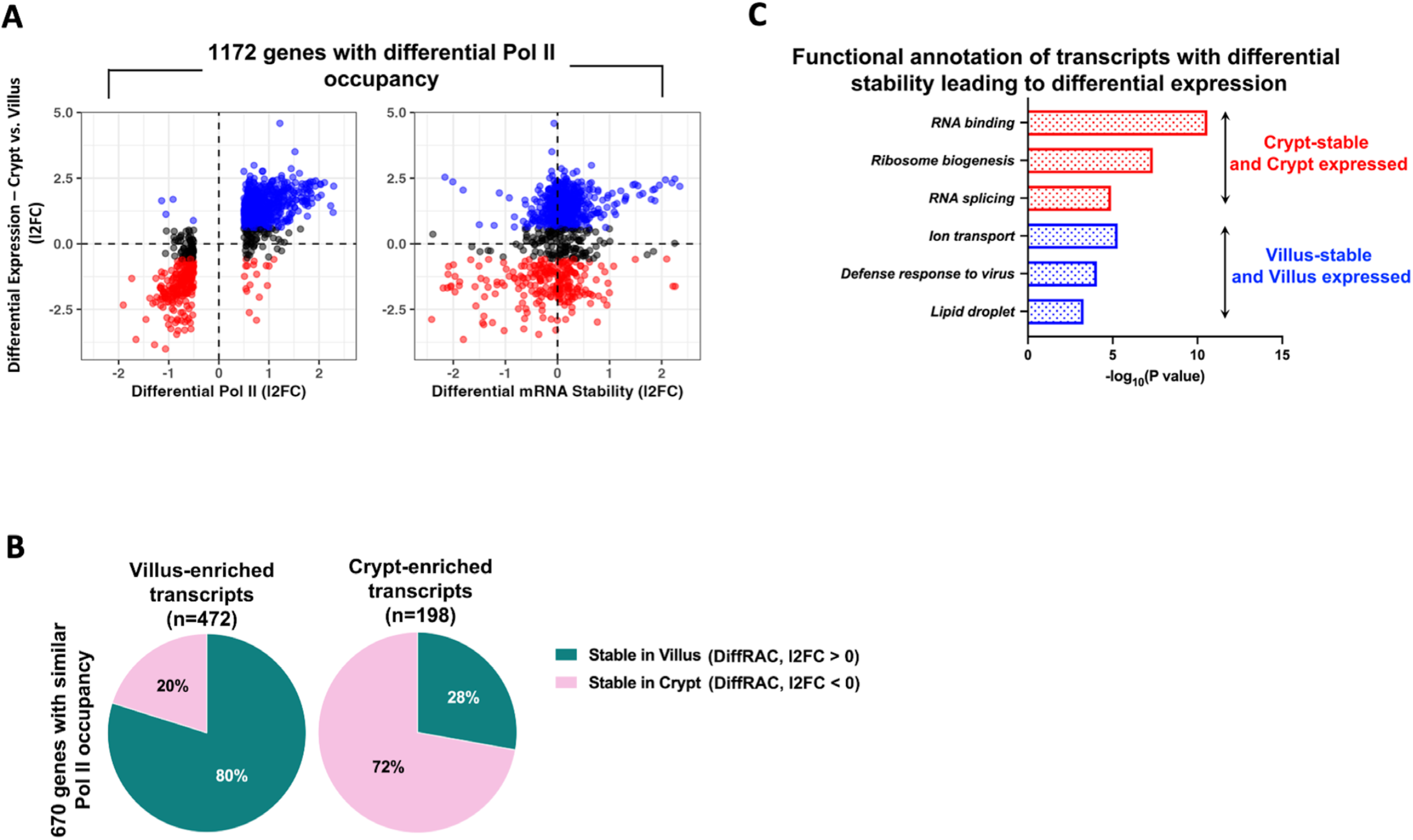
Further characterization of differential mRNA stability analysis. Related to Fig. 2. **(A)** Plot shows 1172 genes with differential Pol II occupancy are less likely to be regulated by differential mRNA stability, as evidenced by their comparison with gene expression patterns in crypt and villus cells (crypt vs. villus RNA-seq: DESeq2 l2FC > 1 or < -1, FDR < 0.05, FPKM > 1; GEO: GSE133949) (see Table S3). **(B)** Analysis of 670 genes showing Pol II occupancy, but differential gene expression reveals that within the group of 472 villus-enriched genes, 80% display stability in villus cells. Conversely, among the 198 crypt-enriched genes, 72% exhibit stability within the crypts (crypt vs. villus RNA-seq and DiffRAC DESeq2, l2FC > 0 or < 0). **(C)** Functional annotation of transcripts with differential stability leading to differential expression. Red bars represent transcripts with greater mRNA stability and higher gene expression levels in the crypts, while blue bars represent transcripts with enhanced mRNA stability and higher gene expression in the villus. p values were calculated using DAVID (see full table in Table S2).

**Fig. S4.**
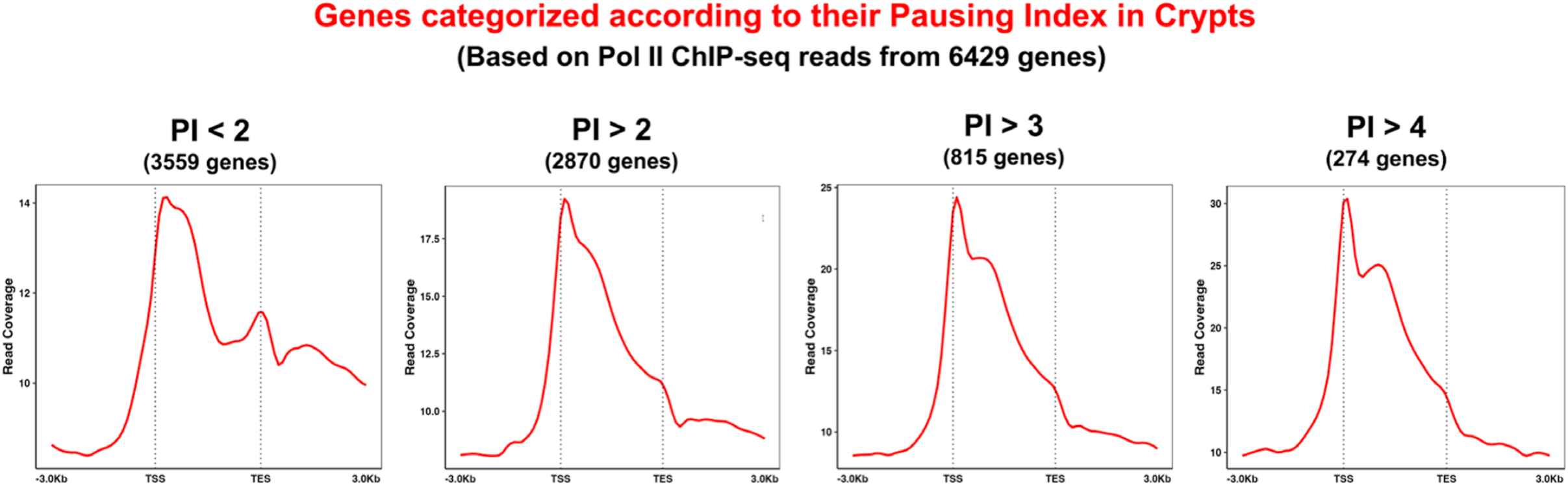
Validation of Pausing Index as a metric for study of promoter-proximal pausing of Pol II. Related to Fig. 3. Pausing indices were calculated in crypt cells and averaged over 3 biological replicates. Genes characterized by a PI > 2 exhibited a clear and distinct peak in Pol II signal at the TSS, which gradually decreased towards the TES. This TSS peak became notably sharper for genes with PI values exceeding 3 and 4. In contrast, genes with a PI less than 2 lacked a discernible TSS peak and typically displayed a higher Pol II signal within the gene body, along with a clustering of Pol II toward the TES.

**Fig.S5.**
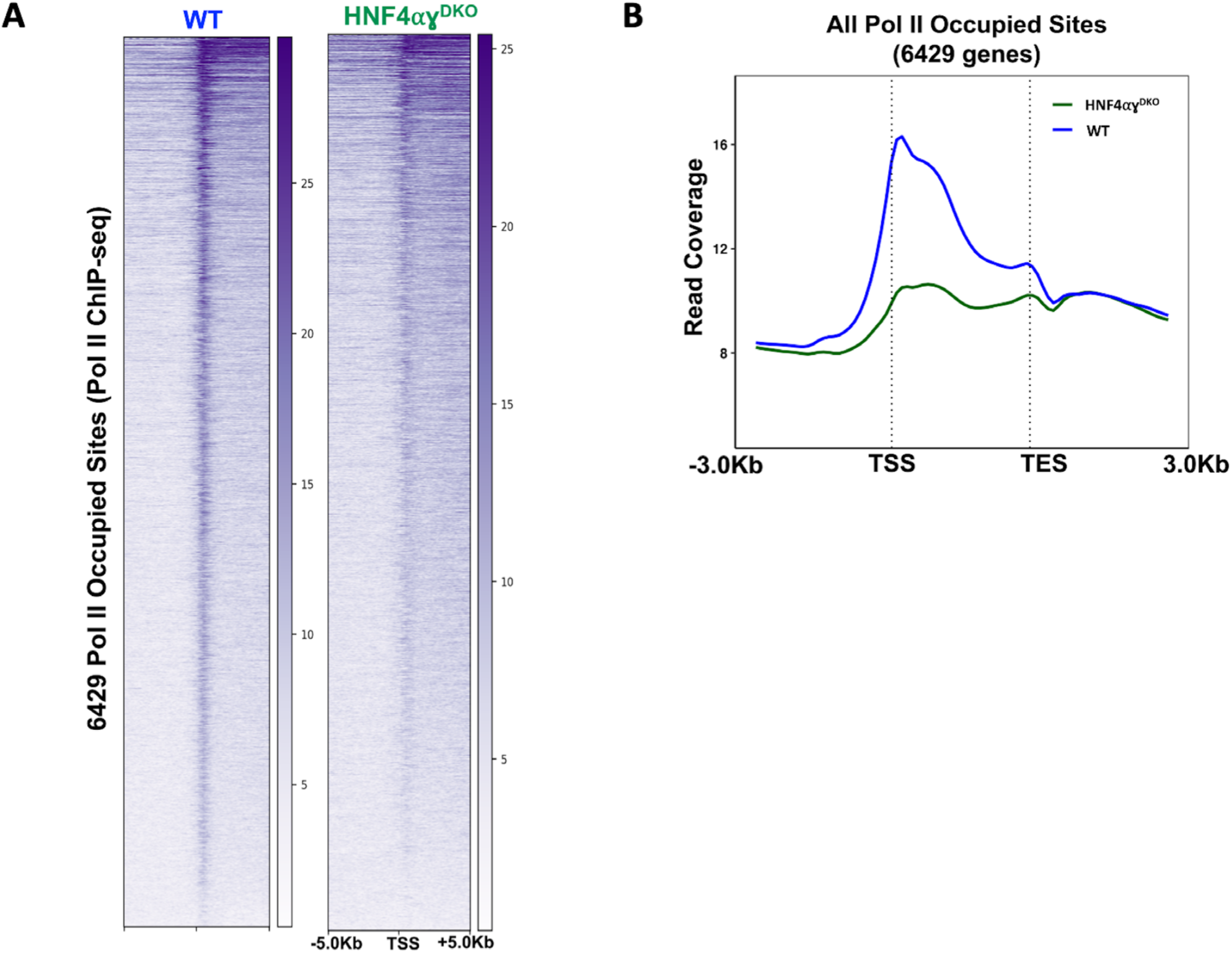
Loss of HNF4 influences Pol II recruitment and gene expression. Related to Fig. 5. **(A)** Heatmap shows reduced binding of Pol II at the promoter regions of 6429 genes (Identified in Fig.S1) in WT and *Hnf4αγ^DKO^* villus cells (Pol II ChIP-seq). **(B)** Metagene profiles of 6429 genes show an overall reduction in Pol II occupancy in *Hnf4αγ^DKO^* cells (Pol II ChIP-seq).

**Fig. S6.**
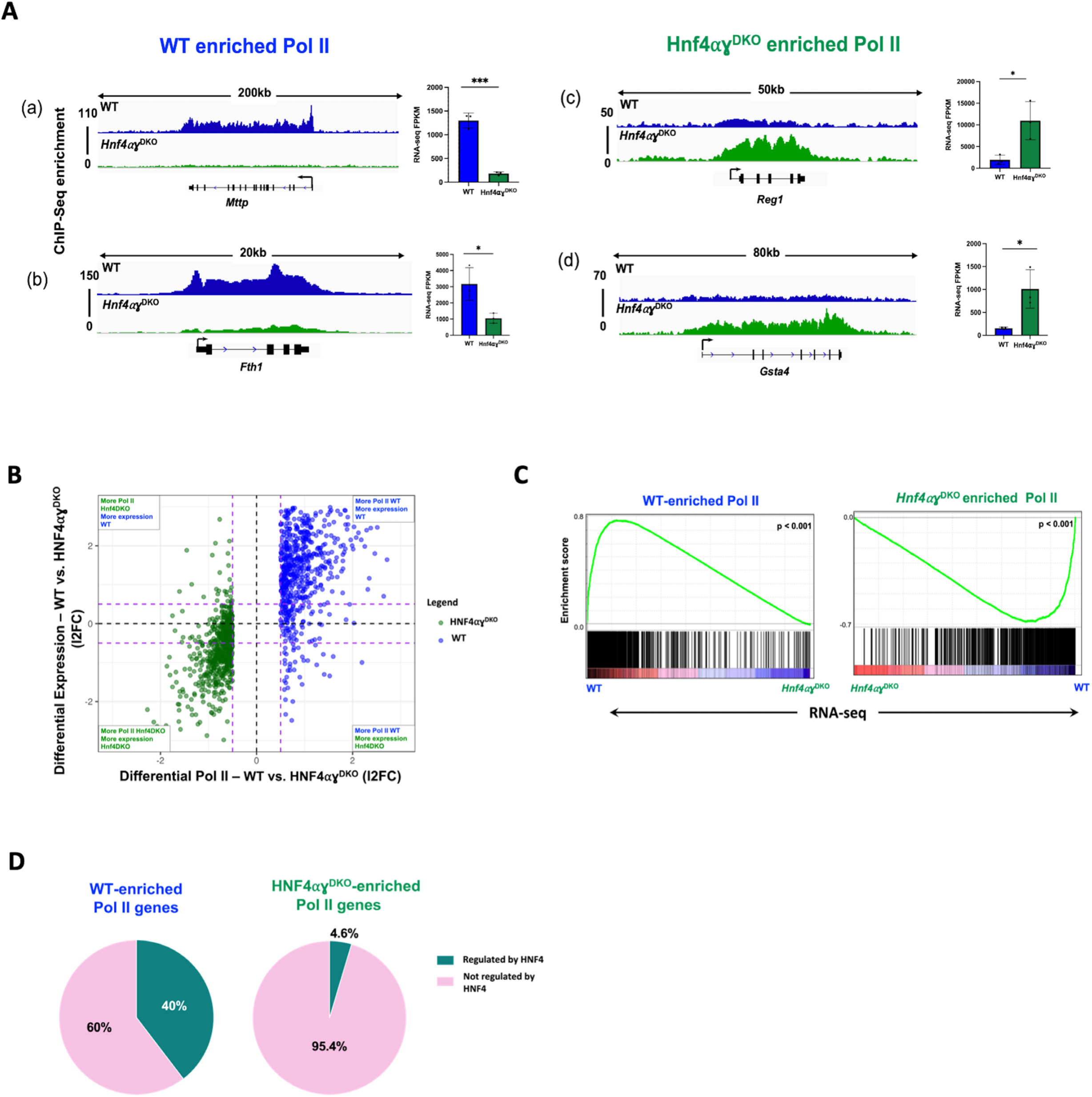
HNF4 promotes Pol II recruitment at its target genes. Related to Fig. 6. **(A)** Representative examples of genes exhibiting WT or *Hnf4αγ^DKO^*-enriched Pol II in villus cells (WT-enriched Pol II genes shown in a-b; *Hnf4αγ^DKO^*-enriched Pol II genes shown in c-d). Loci are as indicated above, data visualized with IGV. WT tracks are depicted in blue and *Hnf4αγ^DKO^* tracks are depicted in green. Bar plots show FPKM values for each gene in WT and *Hnf4αγ^DKO^* cells. FPKM values for each gene derived from WT vs. *Hnf4αγ^DKO^* RNA-seq (GEO: GSE112946). FPKM data is presented as mean ± s.e.m. (n = 3 biological replicates per group). **(B)** Quadrant plots comparing Pol II-enriched genes with WT vs. *Hnf4αγ^DKO^* differential expression patterns. The blue and green points indicate WT- and *Hnf4αγ^DKO^*-enriched Pol II gene sets respectively (differential Pol II: DESeq2, l2FC > 0.58 or < -0.58, FDR < 0.05), which display large magnitude fold changes in Pol II occupancy and differential expression (WT vs. *Hnf4αγ^DKO^* RNA-seq: DESeq2 l2FC > 1 or < -1, FDR < 0.05, FPKM > 1; GEO: GSE112946). The dotted purple lines show a fold change cut-off of 1.5. **(C)** Gene set enrichment analysis (GSEA) reveals that WT-enriched Pol II genes are highly expressed in WT cells, whereas *Hnf4αγ^DKO^* -enriched Pol II genes are highly expressed in *Hnf4αγ^DKO^*cells (Kolmogorov-Smirnov test, p < 0.001). **(D)** Proportion of Pol II enriched genes in WT and *Hnf4αγ^DKO^* - enriched Pol II genes which are dependent on HNF4 (defined as bound and regulated by HNF4 as measured by HNF4A/G ChIP-seq: within 30 kb of HNF4 binding sites; and differentially regulated in WT vs. *Hnf4αγ^DKO^* RNA-seq; GEO: GSE112946). In WT villus cells, 40% of Pol II-enriched genes are HNF4-dependent, while in *Hnf4αγ^DKO^*, only 4.8% of Pol II-enriched genes require HNF4.

## SUPPLEMENTAL TABLES

**Table S1.** Pol II ChIP-seq counts, coordinates and calculated pausing indices from villus/crypt and WT/ *Hnf4αγ^DKO^* cells.

**Table S2.** Gene lists and Gene Ontology analysis (DAVID) of villus/crypt and WT/*Hnf4αγ^DKO^* enriched genes.

**Table S3.** Results of DESeq2 and DiffRAC analyses performed in this publication.

**Table S4.** Results of HOMER *de novo* motif analysis of enhancer sites of genes with villus-enriched gene regulatory events (chromatin looping, Pol II recruitment and gene expression), related to Fig. 4.

**Table S5.** Results of RIME analysis using anti-HNF4α antibodies in primary mouse epithelium, related to Fig. 6.

